# Identifying candidate *de novo* genes expressed in the somatic female reproductive tract of *Drosophila melanogaster*

**DOI:** 10.1101/2023.05.03.539262

**Authors:** Kaelina D. Lombardo, Hayley K. Sheehy, Julie M. Cridland, David J. Begun

## Abstract

Most eukaryotic genes have been vertically transmitted to the present from distant ancestors. However, variable gene number across species indicates that gene gain and loss also occurs. While new genes typically originate as products of duplications and rearrangements of pre-existing genes, putative *de novo* genes - genes born out of previously non-genic sequence - have been identified. Previous studies of *de novo* genes in *Drosophila* have provided evidence that expression in male reproductive tissues is common. However, no studies have focused on female reproductive tissues. Here we begin addressing this gap in the literature by analyzing the transcriptomes of three female reproductive tract organs (spermatheca, seminal receptacle, and parovaria) in three species - our focal species, *D. melanogaster* - and two closely related species, *D. simulans* and *D. yakuba*, with the goal of identifying putative *D. melanogaster*-specific *de novo* genes expressed in these tissues. We discovered several candidate genes, which, consistent with the literature, tend to be short, simple, and lowly expressed. We also find evidence that some of these genes are expressed in other *D. melanogaster* tissues and both sexes. The relatively small number of candidate genes discovered here is similar to that observed in the accessory gland, but substantially fewer than that observed in the testis.

## INTRODUCTION

While the majority of new genes arise through various forms of gene duplication (Long *et al*., 2003), new genes may also arise from ancestrally non-genic DNA (Begun *et al*., 2006; Levine *et al*., 2006). Here we define these *de novo* genes as sequences producing transcripts that are located in ancestrally intergenic DNA and for which there is no evidence of transcription in outgroups. Such transcripts may be coding or non-coding. While putative *de novo* genes have been found in a variety of taxa, including *Drosophila* (*Begun et al., 2006, 2007; Levine et al., 2006; Zhou, Meng, et al., 2008)*, fish (Baalsrud *et al*., 2018; Zhuang & Cheng, 2021), rodents (Casola, 2018; Heinen *et al*., 2009; Murphy & McLysaght, 2012; Neme & Tautz, 2013), plants (Jin *et al*., 2021; Zhang *et al*., 2019) and fungi (Cai *et al*., 2008; Carvunis *et al*., 2012; D. Li *et al*., 2010; Vakirlis *et al*., 2018), our understanding of their possible evolutionary and functional importance remains rudimentary.

Early investigations of *de novo* genes in *Drosophila* provided circumstantial evidence that they may often be expressed in male reproductive tract tissues (Begun *et al*. 2006, Levine *et al*. 2006). That conclusion was supported by subsequent work in *D. melanogaster* (Heames *et al*., 2020; Zhao *et al*., 2014; Zhou, Zhang, *et al*., 2008). Orphan genes found in the *obscura* group of *Drosophila*, some of which may have *de novo* origin, were more likely to be retained if they were highly expressed and male-biased (Palmieri *et al*., 2014). Population level analysis of testis-expressed candidate *de novo* genes in *D. melanogaster* found a total of 142 segregating and 106 fixed genes (Zhao *et al*., 2014). While more conservative criteria reduced the number of candidates (Cridland *et al*., 2022), there were still over 100 polymorphic and 50 fixed genes. A recent investigation of candidate *de novo* genes expressed in the accessory gland (AG; a somatic male reproductive tissue) of *D. melanogaster* revealed 133 candidates, though the AG-expressed genes tended to be fewer in number and expressed less consistently across genotypes than testis-expressed *de novo* gene candidates from the same genotypes (Cridland *et al*., 2022).

While the genetic and population level phenomena that might promote or facilitate expression of candidate *de novo* genes in male reproductive tract tissues remain obscure, the coevolutionary interactions between male and female reproduction raise interesting questions about whether the evolution of male-expressed genetic novelties is correlated with similar processes operating in female reproductive tract (FRT) transcriptomes. Here we begin addressing this question in an analysis of three somatic tissues of the *Drosophila* female reproductive tract, the parovaria, the seminal receptacle, and the spermatheca.

These tissues are all crucial to female reproduction, including ovulation, fertilization, and sperm storage. The parovaria, or female accessory glands, have secretory functions required for fertilization and ovulation (Sun & Spradling, 2012). The seminal receptacle is responsible for short term sperm storage (Fowler, 1973); given the *D. melanogaster* mating system, the seminal receptacle may contain the sperm of multiple males (Manier *et al*., 2010). Spermathecae also serve as sperm storage organs, but are biased toward long-term storage specifically (Pitnick *et al*., 1999). The spermathecae also have secretory cells that participate in fertilization and ovulation pathways (Schnakenberg *et al*., 2011; Sun & Spradling, 2013).

We chose these tissues because the potential for direct interaction of male products, including those produced by novel genes, with the female reproductive tract might generate selection favoring female-expressed genetic novelties. Because these female organs are small and poorly studied, and because *de novo* genes tend to be expressed at low levels, they are unlikely to have been discovered in previous work based on annotations or transcriptome analysis of whole animals or bulk female reproductive tracts. To begin to identify these potentially overlooked *de novo* genes we characterized the transcriptomes of parovaria, spermatheca, and seminal receptacle from mated females from our focal species, *D. melanogaster*, and from two closely related species, *D. simulans*, and *D. yakuba*. Using these data in combination with existing genomic resources, we identified putative *D. melanogaster de novo* genes expressed in these tissues and compared their attributes to those of candidate *de novo* genes previously identified from the study of male *D. melanogaster* reproductive tissues.

## MATERIALS AND METHODS

### Fly strains, datasets used, sequencing and data processing

*Drosophila melanogaster* inbred strains from Raleigh, NC ((Mackay *et al*., 2012); RAL 304, 307, 360, 399, and 517) were raised at 25°C in a 12:12 light:dark cycle. In addition to these inbred lines, an F1 female genotype derived from crossing RAL 304 females x RAL 307 males was also used. Individual 3-5 day old virgin females from lines RAL 304, RAL 307, RAL 360, RAL 399, or RAL 304/307 heterozygotes, were placed in a vial with two RAL 517 males. For *D. simulans* and *D. yakuba*, we used lines Lara 10 (Sedghifar *et al*., 2016) and Tai18E2 (Begun *et al*. 2007), respectively, with females and males being derived from the same strain. Vials were observed, and the time of copulation recorded.

Males were removed immediately after mating, with female reproductive tract tissues dissected within 3-5 hours after the end of copulation, by which time the sperm storage organs are expected to be full (Fowler 1973). The dissected tissues were the parovaria, spermatheca, and seminal receptacle. Dissections were carried out one genotype at a time in a nine-well (3 × 3) depression plate. Wells were filled with roughly 1 ml ice-cold PBS. The initial dissection occurred in the middle well. First, the parovaria was removed from the rest of the reproductive tract and then transferred to a new well, where extra fat or other connecting tissue was removed; the parovaria were then moved to Trizol on ice. Forceps were checked for tissue contamination under the dissecting scope and then rinsed in ethanol. The spermathecae were then removed, rinsed in PBS, and transferred to Trizol on ice. Finally, the seminal receptacle was removed, rinsed in PBS, and then transferred to Trizol on ice. Dissections from roughly ten females were used for each organ and genotype. RNA was extracted from tissues using Trizol, and a cDNA library was produced using a SMART-Seq(R) v4 Ultra(R) Low Input RNA Kit for Sequencing (cat. 634896). Paired-end libraries were generated using the Nextera XT DNA Library Preparation Kit (FC-131-1024) and the Nextera XT Index Kit (FX-131-1001). These libraries were sequenced using 150 bp paired-end reads on an Illumina HiSeq4000 machine at the UC Davis Genome Center.

Trinity (Grabherr *et al*., 2011) was used to assemble transcripts from all reads, left and right, in FASTQ format, from each library. Reads from a total of 12 *D. melanogaster* libraries distributed across five genotypes and three tissues (not all genotypes were assayed for all three tissues) were pooled together and assembled (Supplemental Table 3). Default parameters were used. Raw reads were not trimmed prior to assembly, but instead, rigid downstream filtering (described below) was used to remove poor quality or repetitive sequence. This process aims to retain as much potentially useful data as possible. BLAST was also used to ensure that adapters had been properly removed from candidate *de novo* genes by searching for Illumina Nextera adapter sequences. Several candidate gene transcripts had adapter sequences appended to one end; these were manually trimmed. Trinity was also used at default parameters to assemble transcripts from RNA-seq reads from eight tissue types (whole organism, reproductive tract, gonad, terminalia, thorax, abdomen, viscera, and head) from each of both sexes for eight *Drosophila* species: *D. yakuba, D. ananassae, D. pseudoobscura, D. persimilis, D. willistoni, D. mojavensis, D. virilis*, and *D. grimshawi* (Yang *et al*., 2018). Additional details regarding these samples and reads can be accessed at Gene Expression Omnibus (Edgar *et al*., 2002) under accession numbers GSE99574 and GSE80124. The resulting Trinity transcripts were used to create BLAST databases used to investigate the possibility that outgroup transcripts were homologous to *D. melanogaster de novo* gene candidates, thus falsifying the *de novo* gene hypothesis (see below).

### Criteria for identifying *de novo* gene candidates

While a small fraction of annotated *D. melanogaster* genes might have *de novo* origins, here we focus on currently unannotated candidate genes, as the properties of *de novo* genes - low expression levels, small size, and by definition, absence from related species, makes them much less likely to be annotated. The procedure for identifying candidate *de novo* genes closely follows Cridland *et al*. (2022). Each of the Perl scripts utilized in the pipeline (Supplemental Figure 1), as well as several intermediate steps that called external software or bash procedures, were called from a Python wrapper. Individual Perl scripts and Python wrapper are housed in a GitHub repository (https://github.com/kaelom/Dmel_DNG_Pipeline_2023).

All assembled transcripts were passed through a series of filters. First, we sought evidence for homology between our assembled transcripts and existing annotations or transcripts. To do this, we used BLAST (v. 2.10.1+, Altschul *et al*. 1990) to identify matches to existing annotations against reference genomes from *D. melanogaster* (v. 6.41), *D. yakuba* (v. 1.05), *D. simulans* (v. 2.02), and *D. ananassae* (v. 1.06) from the published Flybase reference (Gramates *et al*., 2022). Corresponding FASTA files contained records for 3’UTR, 5’UTR, intergenic, intronic, miRNA, miscRNA, ncRNA, pseudogene, transposon, tRNA, and CDS (Flybase; http://ftp.flybase.net/releases/FB2020_04/, all downloaded July 23, 2020, with the exception of *D. melanogaster*, http://ftp.flybase.net/releases/FB2021_04/dmel_r6.41/, downloaded August 16, 2021; (Thurmond *et al*., 2019)). Databases created from these resources include: *D. melanogaster* chromosome and intergenic sequence, and the 3’-UTR, 5’-UTR, intronic, miRNA, miscRNA, ncRNA, pseudogene, transposon, tRNA, and CDS for every species available (see Supplemental Table 6 for complete list). In all cases a BLAST match was defined as 80% identity over at least 100 bp. Transcripts that matched existing annotations in any of the four species were removed from further consideration.

To reduce the probability that the surviving *D. melanogaster* transcripts correspond to unannotated outgroup transcripts, we then compared them to a database of *de novo* assembled transcripts created from RNA-seq reads derived from eight tissues (whole organism, gonad, reproductive tract, terminalia, thorax, viscera, head, abdomen) from both sexes from each of the eight aforementioned *Drosophila* species (Yang *et* al., 2018). All unannotated *D. melanogaster* FRT transcripts that returned a BLAST match, as defined above, to any of these resources, were removed.

The remaining *D. melanogaster* candidates were retained only if the transcript sequence was > 300 bp long, the distance to the nearest exon annotation was > 250 bp, and were intergenic (did not reside within annotated *D. melanogaster* introns). Additionally, candidates were required to be expressed at TPM ≥ 1 in at least one FRT library. TPMs were estimated using HISAT2 and StringTie (v2.2.1 (Kim *et al*., 2019); v2.1.4, (Pertea *et al*., 2015)). First, HISAT2 was used to produce SAM files for each library by aligning our FRT raw reads with databases made from the *D. melanogaster* reference genome (v. 6.41) which were then converted to sorted BAM files with Samtools (v1.9, (H. Li *et al*., 2009). These BAM files, along with a *D. melanogaster* reference GTF (Flybase; http://ftp.flybase.net/releases/FB2020_04/dmel_r6.35/gtf/;downloaded August 16, 2021), updated to include Cridland *et al*. (2022) AG candidates as well as our FRT *de novo* candidates, were used to create new GTF and abundance files for each library with StringTie, resulting in species and tissue expression estimates for each candidate transcript. For candidates with multiple isoforms, all isoforms were retained as long as all of these criteria were met for at least one isoform.

Finally, to provide further support for *de novo* origination and reduce the probability that incomplete and/or erroneous genome assemblies lead to errors in *de novo* gene identification, we performed a microsynteny analysis so that sequence homologous to the region corresponding to a *de novo* gene candidate could be identified in the orthologous regions of *D. simulans* and *D. yakuba*. To do so, we identified the annotated genes immediately upstream or downstream of each candidate. Then, the Flybase 2021 ortholog database (Flybase; http://ftp.flybase.net/releases/FB2021_02/precomputed_files/orthologs/; downloaded May 12, 2021) was used to identify the orthologs in the outgroups. A FASTA file containing those genes, the candidate, and 5 kb downstream and upstream of the region was then produced. BLAST analysis of these regions was then performed to identify these micro-syntenic regions in the reference chromosome databases for *D. simulans* and *D. yakuba*. A file was produced that contained the positions of orthologous matches, if they existed. Because of the small number of candidate genes identified here, we retain as weaker candidates those genes that failed this final synteny step with *D. simulans* or *D. yakuba*. Candidates that did not show positive evidence of syntenic regions in one or both outgroups were checked manually over larger genomic regions in the UCSC GenomeBrowser (Karolchik *et al*., 2003).

### Expression in existing transcriptome resources

We investigated expression of our candidate genes in data from a previously published analysis of *D. melanogaster* female reproductive tract transcriptomes (McDonough-Goldstein *et al*., 2021), derived from six tissues from unmated and mated females: bursa, oviduct, seminal receptacle, spermathecae, parovaria, and the FRT-associated fat body. Tissues from mated females were collected 6h and 24h post-mating. The *D. melanogaster* reference GTF, (v6.41) was updated to include our *de novo* gene candidates. We then used StringTie (Pertea *et al*., 2015) with default parameters to estimate TPMs of the identified FRT candidate genes using the reads from McDonough *et al*. 2021. Additionally, we used reads from FlyAtlas2 (Krause *et al*., 2022; Leader *et al*., 2018), using the StringTie methods described above, to estimate expression of our candidate genes in a variety of *D. melanogaster* tissues. We used the same approach to investigate expression in our female data of previously identified candidate *de novo* genes expressed in the accessory gland + anterior ejaculatory duct of Raleigh inbred lines (Cridland *et al*. 2022).

### Coding potential

Coding potential of putative *de novo* genes was assessed with two different methods, Coding Potential Calculator 2 (CPC2) and Coding Potential Assessment Tool (CPAT) (Kang *et al*., 2017; Wang *et al*., 2013). CPAT provides specific default parameters depending on the query species, therefore the default parameters for *Drosophila* were used. Settings for CPC2 are not dependent on the species being investigated. Browser versions of each tool were used at default parameters. CPAT and CPC2 each had their own proprietary coding potential cutoff of 0.39 and 0.5, respectively. CPAT’s default minimum ORF length is 75 nucleotides, while CPC2 does not enforce a minimum.

Protein translations of these transcripts were run through SignalP 6.0, at default settings, in order to assess candidates for signal sequences (Teufel *et al*., 2022).

## RESULTS

### *De novo* genes identified and basic characteristics

We identified 61 candidate *de novo* transcripts (Supplemental File 1) associated with 35 *de novo* gene candidates (TABLE 1), which were expressed in *D. melanogaster* but for which we found no evidence of expression in *D. simulans* or *D. yakuba*. None of these candidates exhibited evidence of homology with transcripts observed from any tissue in any of the additional seven Drosophila species examined (Methods). Of these 35 candidates, 32 had validated microsynteny with *D. simulans*, and 29 of those also had confirmed microsynteny with *D. yakuba*.

**Table 1.** Summarized Candidate Expression Across Datasets and Tissues

| ID | Number of libraries expressed this report | Number of tissue x mating status expressed McDonough et al. | Number of tissues expressed FlyAtlas2 | FRT Tissue Expression (This report, **this report** and McDonough et al., McDonough et al.) | FlyAtlas2 Average TPMs |  |  | Max TPM |
| --- | --- | --- | --- | --- | --- | --- | --- | --- |
|  |  |  |  |  | Male | Female | 3rd Instar Larvae |  |
| TRINITY_DN18465_c0_g4 | 5 | 2 | 15 | **SR, ST, Bursa** | Brain (7.11), Crop (1.46), Eye (4.80), Head (2.72), TG (5.30) | Brain (7.00), Crop (1.36), Eye (4.41), Head (2.95), Ovary (1.46), TG (5.56), Whole (1.49), Carcass (1.02) | Larval CNS (9.86), Larval Trachea (1.56) | 2.05 |
| TRINITY_DN58663_c0_g1 ** | 1 | 10 | 4 | **SR, ST, Bursa, Oviduct** | Whole (1.40), MG (1.00), TG (1.10) | TG (1.40) | - | 1.06 |
| TRINITY_DN2173_c0_g1 | 3 | 3 | 2 | **ST** | Eye (3.72) | Eye (1.07) | - | 7.00 |
| TRINITY_DN52147_c0_g1 | 4 | 3 | - | **SR** | - | - | - | 1.24 |
| TRINITY_DN90_c0_g3 | 5 | - | - | ST, PV | - | - | - | 1.59 |
| TRINITY_DN16805_c0_g2 | 2 | 3 | - | **ST** | - | - | - | 1.81 |
| TRINITY_DN65427_c0_g1 | 1 | - | 3 | ST | Brain (2.22) | Brain (1.66), TG (1.21) | - | 1.09 |
| TRINITY_DN40913_c0_g1 | 1 | - | 3 | ST | Brain (1.22), TG (2.89) | TG (2.29) | - | 1.23 |
| TRINITY_DN7862_c0_g3 | 3 | - | - | SR | - | - | - | 3.29 |
| TRINITY_DN5611_c0_g1 | 2 | 1 | - | **ST** | - | - | - | 2.47 |
| TRINITY_DN4094_c0_g1 ** | 2 | 1 | - | **SR** | - | - | - | 1.09 |
| TRINITY_DN66768_c0_g1 | 1 | - | 2 | ST | Anal Pad (6.27) | Anal Pad (5.38) | - | 1.43 |
| TRINITY_DN7830_c0_g1 | 1 | - | 2 | ST | Anal Pad (3.93) | Anal Pad (8.41) | - | 3.19 |
| TRINITY_DN2143_c0_g2 | 1 | - | 2 | ST | - | VST (1.81), MST (1.69) | - | 1.63 |
| TRINITY_DN3265_c1_g1 | 1 | 2 | - | **SR** | - | - | - | 1.26 |
| TRINITY_DN72444_c0_g1 | 2 | - | - | ST | - | - | - | 1.68 |
| TRINITY_DN7278_c0_g1 | 1 | - | 1 | ST | AG (1.96) | - | - | 1.07 |
| TRINITY_DN24046_c0_g1 | 1 | - | 1 | PV | - | Brain (1.03) | - | 1.21 |
| TRINITY_DN4410_c0_g1 | 2 | - | - | SR, ST | - | - | - | 1.22 |
| TRINITY_DN4173_c1_g1 | 2 | - | - | SR | - | - | - | 1.50 |
| TRINITY_DN11963_c0_g1 | 1 | - | 1 | ST | Hindgut (1.18) | - | - | 1.59 |
| TRINITY_DN47998_c0_g1 | 2 | - | - | SR | - | - | - | 1.99 |
| TRINITY_DN6533_c0_g1 * | 2 | - | - | SR | - | - | - | 1.81 |
| TRINITY_DN22183_c0_g1 | 2 | - | - | SR | - | - | - | 1.65 |
| TRINITY_DN41677_c0_g1 | 1 | - | - | PV | - | - | - | 1.01 |
| TRINITY_DN15918_c0_g3 | 1 | - | - | ST | - | - | - | 1.27 |
| TRINITY_DN22384_c0_g1 | 1 | - | - | ST | - | - | - | 1.45 |
| TRINITY_DN13179_c1_g1 | 1 | - | - | ST | - | - | - | 1.40 |
| TRINITY_DN43648_c0_g1 * | 1 | - | - | SR | - | - | - | 1.40 |
| TRINITY_DN27150_c0_g4 | 1 | - | - | SR | - | - | - | 1.25 |
| TRINITY_DN55991_c0_g1 | 1 | - | - | SR | - | - | - | 1.48 |
| TRINITY_DN12518_c0_g2 * | 1 | - | - | SR | - | - | - | 1.03 |
| TRINITY_DN65366_c0_g1 ** | 1 | - | - | SR | - | - | - | 1.20 |
| TRINITY_DN10735_c0_g1 | 1 | - | - | SR | - | - | - | 1.15 |
| TRINITY_DN5533_c0_g1 | 1 | - | - | ST | - | - | - | 1.88 |

\* Indicates that the candidate has microsynteny in *D. simulans* only. \*\* Indicates that the candidate has no microsynteny in either outgroup. FlyAtlas2 - numbers beside tissue label indicate the average TPM for all replicates of that tissue and sex. FRT Expression - It is important to note that for a candidate that was significantly expressed in a certain FRT tissue across our 12 libraries, that candidate was also significantly expressed in only the corresponding tissue(s) in the McDonough et al. FRT dataset when **bolded**. Candidates expressed only in a certain tissue in McDonough et al. are *italicized*.

Most transcripts (44/61, 72%) had a simple structure, containing only one exon, though some contained up to three. Similarly, most candidate genes were associated with a single isoform, though 14 had at least two, and one, TRINITY_DN4410, had as many as seven.

Candidate transcript lengths, considering the longest isoform of each, range from 302 bp to 3477 bp, with average length of 810 bp, similar to the mean length for previously published AG-expressed intergenic candidate genes (701 bp; Cridland *et al*. 2022) and testis-expressed candidate genes (935 bp; Zhao *et al*. 2014). These FRT-expressed candidate genes were distributed roughly homogeneously across chromosome arms (Supplemental Table 1).

### Quantification of expression of candidates in the FRT

Per our filtering criteria, each candidate was expressed at a TPM ≥ 1 in at least one of the 12 tissue/genotype Raleigh inbred line combinations, though several (14/35, 40%) were expressed above this threshold in two to five libraries (Supplemental Table 1). Consistent with previous reports that *de novo* genes tend to be lowly expressed, the mean TPM of expressed candidate genes (including only observations of TPM ≥ 1) was only 1.57. Thus, many candidates were expressed at only slightly higher levels than the minimum TPM criterion for expression and only in one library. The maximum observed TPM was 7 (Table 1, Supplemental Table 1). To further summarize candidate *de novo* gene expression relative to expression levels of annotated genes, we calculated, per library, the number of annotated genes that were expressed over the maximum TPM observed among our candidates for that library. We found that 43% (in the RAL 307 ST library) to 71% (in the RAL 304 PV library) of annotated genes exhibited TPM > candidate *de novo* gene max TPM. All *de novo* gene candidates had non-zero TPMs in multiple libraries, five on average.

### Candidate expression in other female reproductive tract tissues

We took advantage of two existing RNA-seq resources to seek further evidence of expression of our 35 candidate genes. We used a published transcriptome analysis of the *D. melanogaster* female reproductive tract (McDonough *et al*. 2021), which included data from virgin and mated females (two post-copulation time points, six hours and 24 hours) for the same three organs used here, as well as for the bursa, oviduct, and fat body.

In female reproductive tract data from McDonough *et al*. we observed expression (mean TPM across biological replicates for each tissue/mating status ≥ 1) for 8 of 35 (23%) *de novo* gene candidates (Table 1). Maximum TPM of our candidates in any one of these libraries was 5.44. Most TPM ≥ 1 estimates were from either the spermatheca or seminal receptacle, and in many cases, these candidates also showed significant expression in those same tissues in our FRT data (Table 1). Often, expression in these tissues included numerous mating statuses (Supplemental Table 4).

### Expression of candidate *de novo* genes in FlyAtlas 2 data

To investigate evidence of broader candidate *de novo* gene expression in publicly available transcriptome data, we used the RNA-seq reads from FlyAtlas2 (Leader *et al*. 2018), which includes data from several adult male and female tissues, and 3rd instar larval tissues. Expression analysis was as described above, where TPM estimates for replicates and isoforms were pooled and averaged for each candidate. Genes were categorized as expressed in a tissue if mean TPM was greater than or equal to one. Eleven of 35 (31%) FRT-expressed candidates exhibited TPM ≥ 1 in at least one tissue in FlyAtlas2, with the maximum TPM from any one library being 12.63. Ten candidates were expressed in one to four of these tissues, but one candidate, TRINITY_DN18465_c0_g4, was expressed in 15 different tissues (Table 1, Supplemental Table 5).

### Expression of accessory gland expressed *de novo* genes in female reproductive tissues

Two of the organs used in our experiments, the spermatheca and parovaria, contain secretory cells (Allen & Spradling, 2008; Mayhew & Merritt, 2013). Because the male accessory gland is composed primarily of secretory cells (Wilson *et al*., 2017), we investigated whether previously identified accessory gland-expressed *de novo* gene candidates (Cridland *et al*. 2022) are expressed (TPM ≥ 1) in our FRT data. Twelve of the 133 AG-expressed candidates (about 9%) were also expressed in at least one FRT library (Supplemental Table 2). This supports the conclusion from the analysis of the McDonough *et al*. and FlyAtlas2 data that the FRT-expressed candidate *de novo* genes are not always tissue-specific, or even sex-specific in their expression. Whether or not the candidates expressed in FRT are biased toward secretory cell expression is an open question. Two of these twelve AG-expressed *de novo* genes that were also identified from our female tissues, TRINITY_DN4679_c0_g1 and TRINITY_DN42840_c0_g1, were also expressed (TPM ≥ 1) in the FRT data from McDonough *et al*. 2021. Maximum expression of TRINITY_DN4679 in any one library was exhibited in the unmated bursa, 3.36 TPM, while the highest expression of TRINITY_DN42840, TPM = 4.14, was exhibited in the 6hr post-mating oviduct.

Overall, of the 35 candidates pre-microsynteny-validation, 24 (69%) were called as expressed in two or more of our libraries, or at least twice, inclusive of our libraries and previously published RNA-seq datasets (Table 1). Of the 29 candidates genes with confirmed synteny to both outgroups, 12 (41%) were expressed at TPM ≥ 1 in more than one FRT library (Table 1), and 14 (48%) were expressed at mean TPM ≥ 1, in at least one of two previously published RNA-seq datasets (FlyAtlas 2 and McDonough *et al*. 2021; Table 1).

Given the heterogeneous genotypes, experimental conditions, and RNA-seq data production methods across these resources, this should be viewed as a lower bound on the repeatability of candidate gene transcript production in *D. melanogaster*.

### Coding potential

Two methods, CPAT and CPC2, were used to investigate coding potential of candidate transcripts. CPAT analysis revealed that while all 61 candidate transcripts had ORFs that exceeded the length minimum of 30 nucleotides, two transcripts, both isoforms of TRINITY_DN7862_c0_g3_(i1,i2) were identified as potentially coding. Isoform 1 of TRINITY_DN7862_c0_g3 was 428 bp long and single-exon, while isoform 2 was 390 bp long and consisted of two exons. In our data, this gene was expressed only in the seminal receptacle of multiple lines: The RAL 304 (3.29 TPM), RAL 307 (2.94 TPM), and RAL 360 (1.18 TPM) (Supplemental Table 1).

CPC2 identified four candidate transcripts as potentially coding: two isoforms of TRINITY_DN16805_c0_g2_(i1, i3), TRINITY_DN47998_c0_g1_i1 and TRINITY_DN90_c0_g3_i1. TRINITY_DN16805 is a single exon candidate that has three isoforms overall. One of these isoforms is only 250 bp long, however, the other two isoforms, which were the two rated as potentially coding, are 1105 and 1147 bp long. This candidate is expressed at TPM ≥1 in the spermatheca of RAL 307 and RAL 307 x RAL 304 F1 (1.81 TPM, 1.18 TPM). Both other potentially coding candidates are single isoform and single exon. TRINITY_DN47998 has a length of 617 bp and was expressed in two FRT libraries: RAL 307 × 304 F1 seminal receptacle (2 TPM) and RAL 360 seminal receptacle (1.37 TPM). TRINITY_DN90 is 486 bp long and was expressed at TPM ≥ 1 in five of our FRT libraries: RAL 304 spermatheca (1.14 TPM), RAL 304 x RAL 307 F1 parovaria (1.40 TPM), RAL 304 x RAL 307 F1 spermatheca (1.05 TPM), RAL 307 parovaria (1.58 TPM) and RAL 307 spermatheca (1.40 TPM) (Supplemental Table 1).

None of the ORFs associated with any candidates were predicted by SignalP to have a signal sequence, which is unsurprising considering the low number of potentially coding candidates.

## DISCUSSION

Our investigation of putative *de novo* genes expressed in three organs of the *D. melanogaster* female reproductive tract, the parovaria, seminal receptacle, and spermatheca, revealed multiple similarities to candidate *de novo* genes expressed in the accessory gland + anterior ejaculatory duct (Cridland *et al*. 2022) of the same population. For example, the number of intergenic *de novo* gene candidates identified here (n = 35) is comparable to that observed in the accessory gland + anterior ejaculatory duct (n= 49) from a similar sample of the same population, though direct quantitative comparison is difficult due to differences between studies in the number of distinct organs sampled (two in Cridland *et al*. (2022) and three here) and number of genotypes used (six inbred genotypes in Cridland *et al*. 2022 vs. four inbred genotypes and one heterozygous genotype here). Also similar to observations from the intergenic candidates expressed in the accessory gland + ejaculatory duct, none were expressed in all genotypes examined. Further similarities include their short length, simple organization, and low expression levels (Table 1, Supplemental Table 1), features shared more generally by candidate *de novo* genes in multiple taxa (reviewed in Van Oss and Carvunis 2019). While *D. melanogaster* testis-expressed candidate *de novo* genes share similarities with those expressed in the accessory gland and female reproductive tract, including their size, simplicity, and relatively low expression, they are much more abundant, and are expressed in a greater proportion of genotypes (Zhao *et al*. 2014, Cridland *et al*. 2022). Whether the Drosophila female germline exhibits patterns of *de novo* gene expression similar to that of the male germline is an important unanswered question.

The expression of candidate genes in different female reproductive tract organs and in other tissues suggests that they are frequently not tissue- or organ-specific. Several are not sex-limited in expression, as in addition to expression in the parovaria, spermatheca, or seminal receptacle, their transcripts are present in the male accessory gland and multiple tissues in both sexes. Whether this is a general property of *D. melanogaster*-specific *de novo* gene candidates is an open question. While computational analysis of the putative *de novo* genes identified here provides little support that most are protein-coding, firmer conclusions on this point await analysis of proteomic or ribo-profiling data (Zheng & Zhao, 2022), or transgenic analysis of epitope-tagged individual candidates.

Finally, given their generally low expression levels and the fact they are not reliably expressed in all *D. melanogaster* genotypes, it remains to be seen whether genetic analysis will provide robust conclusions regarding the functions of these genes. Recent work on the transcriptional behavior of “naive” human DNA in yeast has suggested that eukaryotic DNA has inherent properties associated with the production of mature mRNAs (Luthra *et al*., 2022). Though the relevance of such observations to the attributes of endogenous intergenic DNA in Drosophila is unclear, this finding has potential relevance to the evolution of *de novo* genes. *De novo* genes, especially those that have low population frequencies, may be enriched for the products of some form of background (i.e., spurious) transcription (Begun *et al*. 2006), and if so, their presence and ability to be properly processed by the cell is not particularly informative about the probability that they have specific biological functions. Second, the properties of ancestrally intergenic DNA must necessarily lead to occasional production of spurious transcripts, a small subset of which may have or acquire functions that drive their spread through populations under directional selection (Begun *et al*. 2006, Zhao *et al*. 2014). Genetic and/or population genetic data (e.g., Zhao *et al*. 2014) will be necessary to elucidate the possible evolutionary and/or biological significance of these very young Drosophila genes.

## DATA AVAILABILITY STATEMENT

Female reproductive tract sequences available at https://www.ncbi.nlm.nih.gov/sra under BioProject accession number PRJNA924827. Pipeline scripts and information can be found at https://github.com/kaelom/Dmel_DNG_Pipeline_2023. Supplemental File 1 contains the trimmed transcripts associated with the putative de novo gene candidates, as described above, in FASTA format.

## ACKNOWLEDGEMENTS

This work was supported by the National Institutes of Health grant NIGMS R35GM134930 to DJB The content is solely the responsibility of the authors and does not necessarily represent the official views of the National Institutes of Health. We thank the UC Davis Genome Center for advice and sequencing.

---

**SUPPLEMENTAL FIGURE 1.**
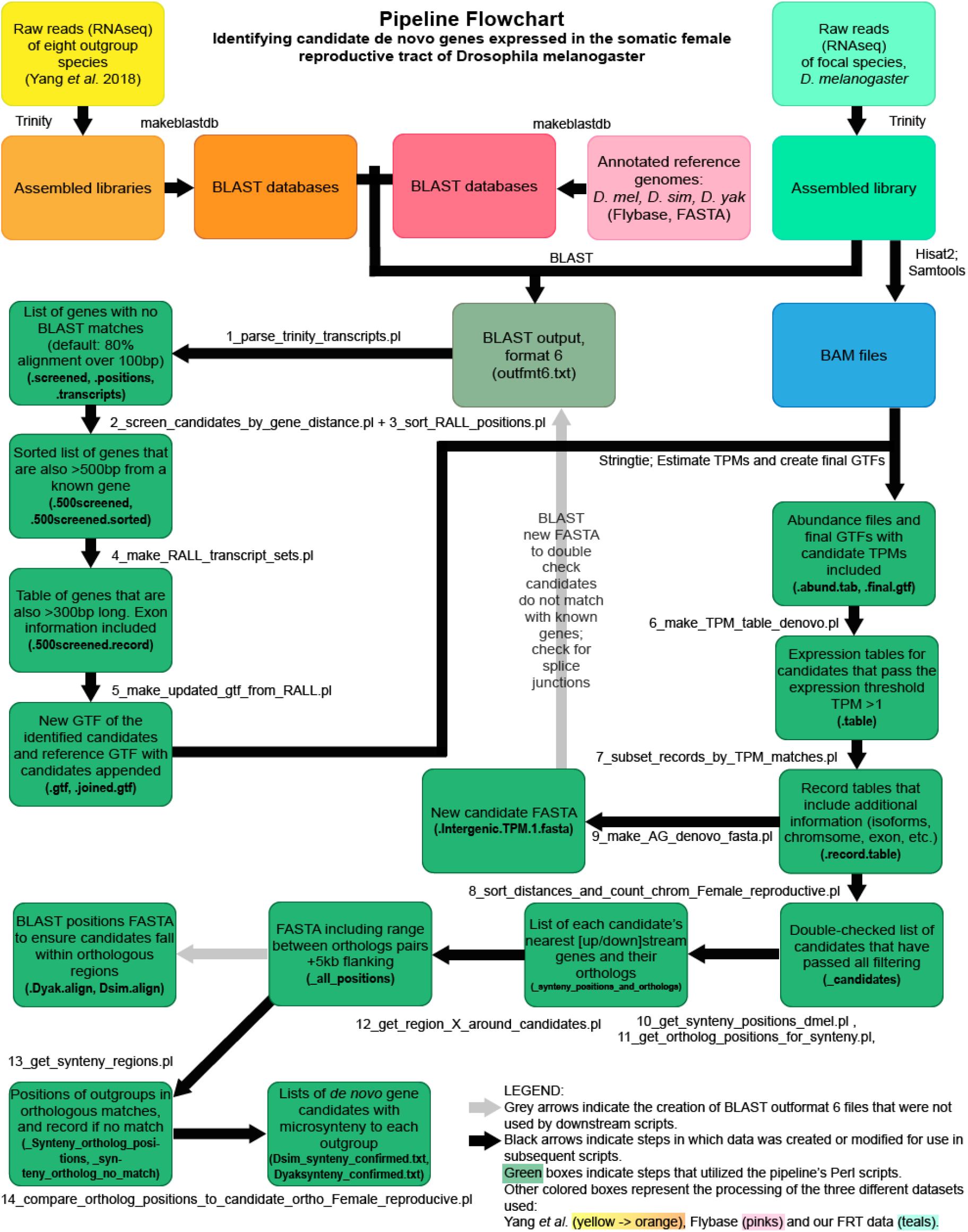
Candidate de novo Gene Identification Pipeline Flowchart.

**Supplemental Table 1.**
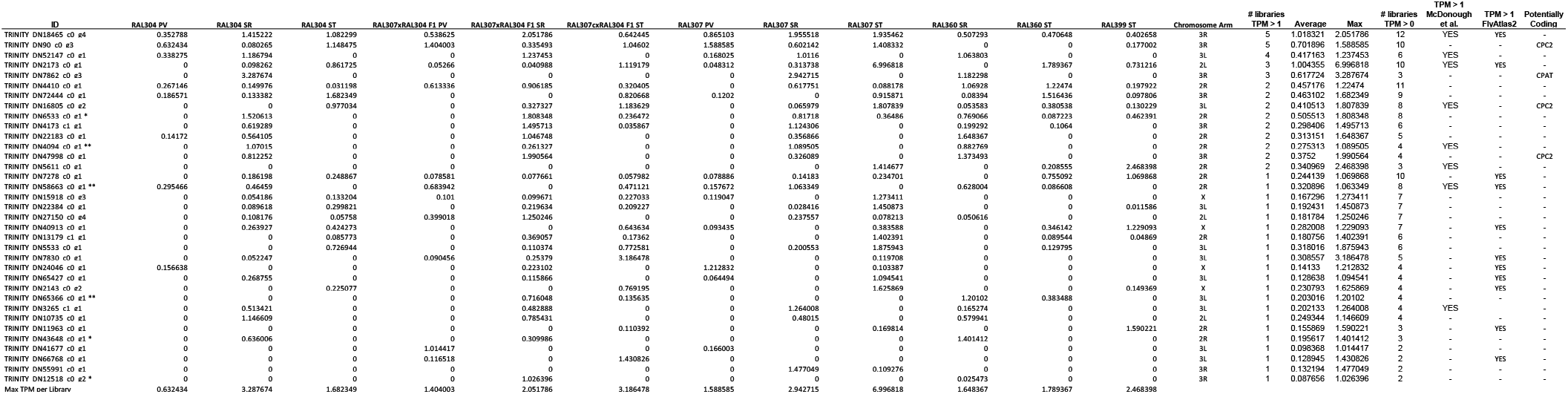
Candidate TPMs in All FRT libraries

**Supplemental Table 2.**
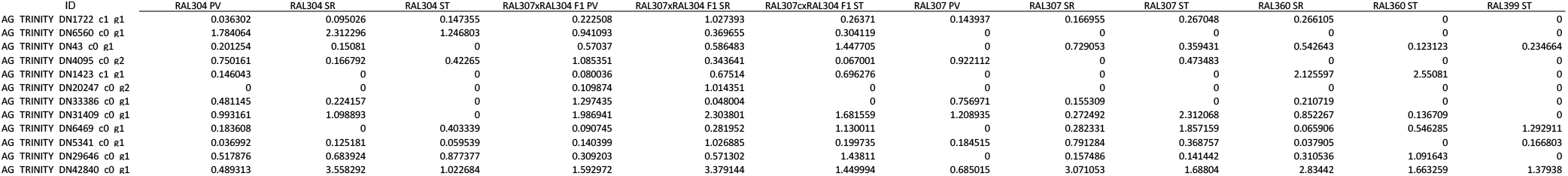
TPMs of Accessory Gland Candidates with TPM ≥ 1 in All FRT Libraries

**Supplemental Table 3.** Raw Read Counts for All FRT Libraries

| Raw Read Files (Left and Right) | Number of Reads |
| --- | --- |
| 304_PV_S195_L003_R1_001.fastq.gz | 20513346 |
| 304_PV_S195_L003_R2_001.fastq.gz | 20513346 |
| 307_PV_S198_L003_R1_001.fastq.gz | 16183938 |
| 307_PV_S198_L003_R2_001.fastq.gz | 16183938 |
| 304_SR_S194_L003_R1_001.fastq.gz | 19797634 |
| 304_SR_S194_L003_R2_001.fastq.gz | 19797634 |
| 307_SR_S197_L003_R1_001.fastq.gz | 14127659 |
| 307_SR_S197_L003_R2_001.fastq.gz | 14127659 |
| 304_ST_S193_L003_R1_001.fastq.gz | 18778304 |
| 304_ST_S193_L003_R2_001.fastq.gz | 18778304 |
| 307_ST_S196_L003_R1_001.fastq.gz | 21357080 |
| 307_ST_S196_L003_R2_001.fastq.gz | 21357080 |
| 360_ST_S199_L003_R1_001.fastq.gz | 18061012 |
| 360_ST_S199_L003_R2_001.fastq.gz | 18061012 |
| 360_SR_S200_L003_R1_001.fastq.gz | 15488884 |
| 360_SR_S200_L003_R2_001.fastq.gz | 15488884 |
| 399_ST_S201_L003_R1_001.fastq.gz | 17549951 |
| 399_ST_S201_L003_R2_001.fastq.gz | 17549951 |
| 307304_PV_S210_L003_R1_001.fastq.gz | 18656859 |
| 307304_PV_S210_L003_R2_001.fastq.gz | 18656859 |
| 307304_SR_S209_L003_R1_001.fastq.gz | 19029027 |
| 307304_SR_S209_L003_R2_001.fastq.gz | 19029027 |
| 307304_ST_S208_L003_R1_001.fastq.gz | 20341929 |
| 307304_ST_S208_L003_R2_001.fastq.gz | 20341929 |
| Tai18_PV_S207_L003_R1_001.fastq.gz | 19049430 |
| Tai18_PV_S207_L003_R2_001.fastq.gz | 19049430 |
| Tai18_SR_S206_L003_R1_001.fastq.gz | 19596835 |
| Tai18_SR_S207_L003_R2_001.fastq.gz | 19596835 |
| Tai18_ST_S205_L003_R1_001.fastq.gz | 21481121 |
| Tai18_ST_S205_L003_R2_001.fastq.gz | 21481121 |
| LARA10_PV_S204_L003_R1_001.fastq.gz | 18161209 |
| LARA10_PV_S204_L003_R2_001.fastq.gz | 18161209 |
| LARA10_SR_S203_L003_R1_001.fastq.gz | 20153219 |
| LARA10_SR_S203_L003_R2_001.fastq.gz | 20153219 |
| LARA10_ST_S202_L003_R1_001.fastq.gz | 23805531 |
| LARA10_ST_S202_L003_R2_001.fastq.gz | 23805531 |

**Supplemental Table 4.**
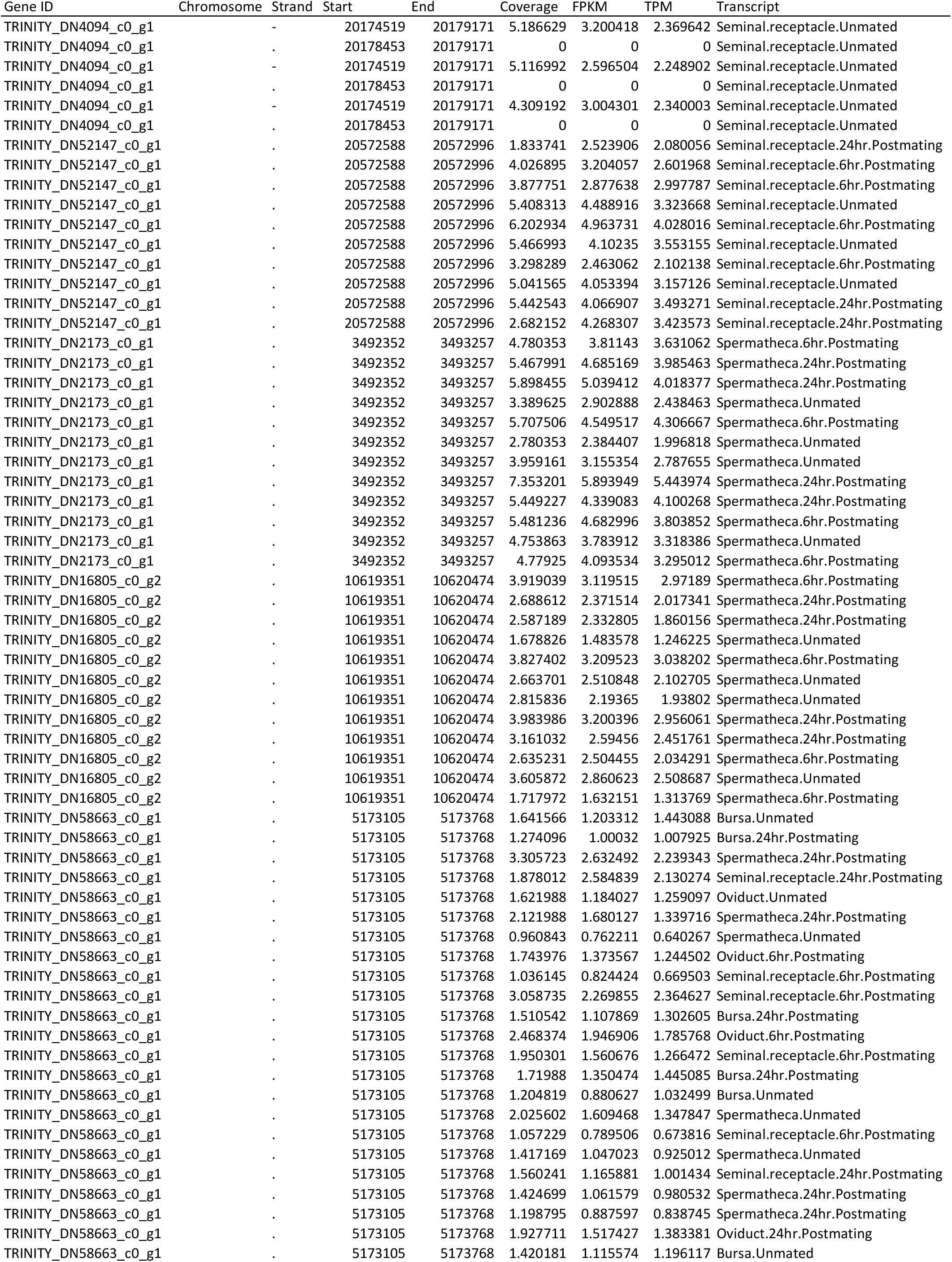
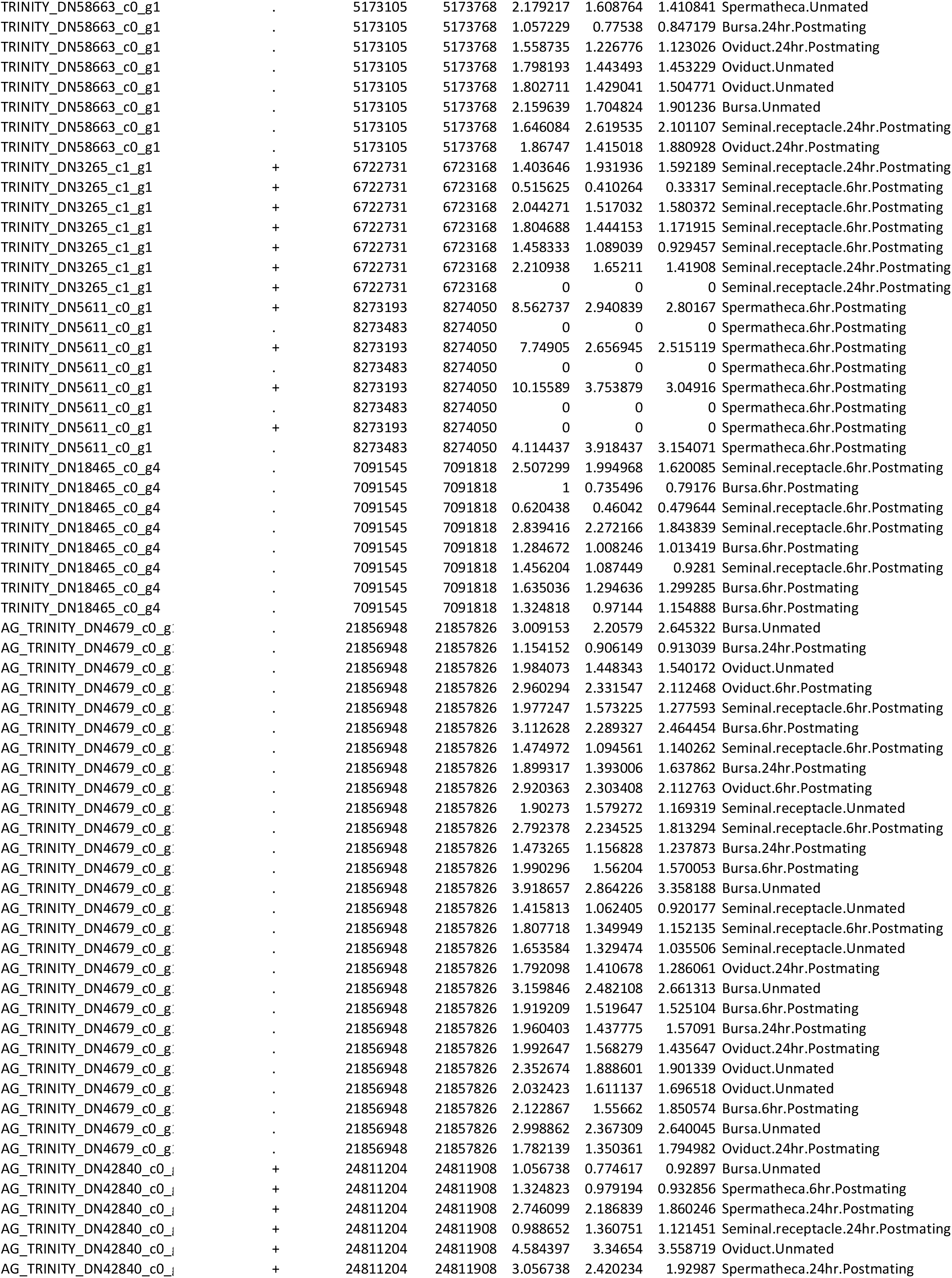
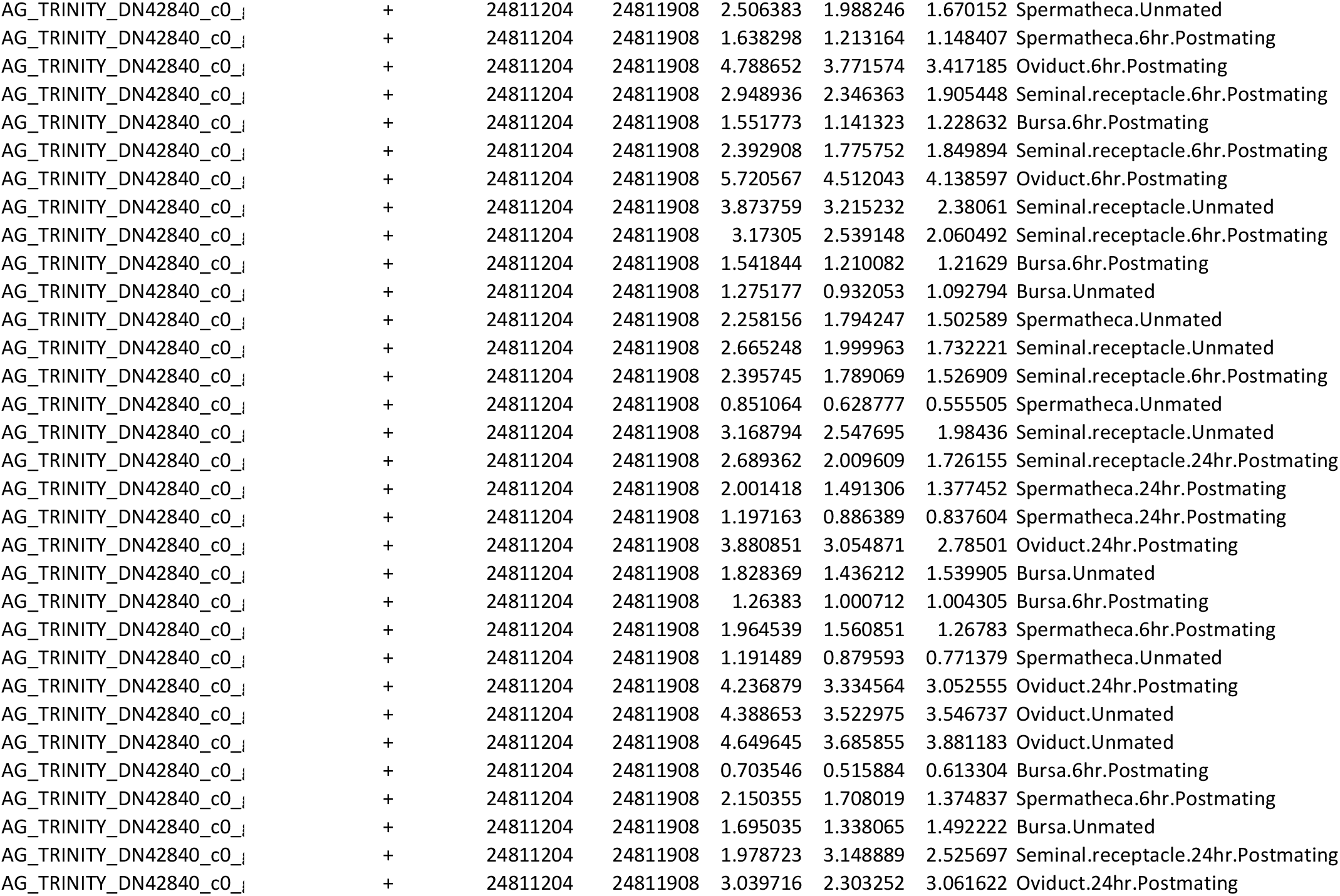
Expressed Candidate TPMs in McDonough *et al*. Libraries

**Supplemental Table 5.**
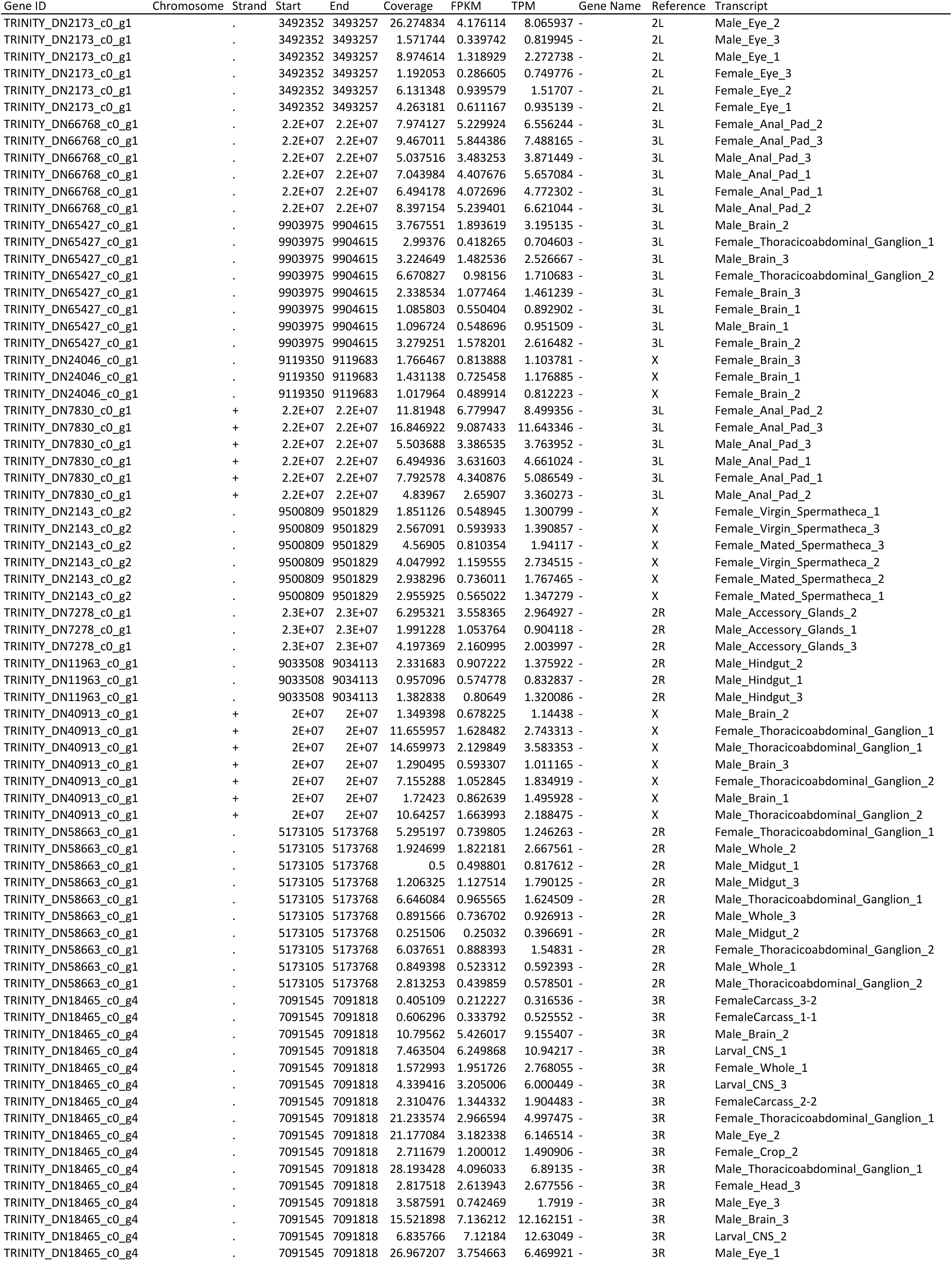
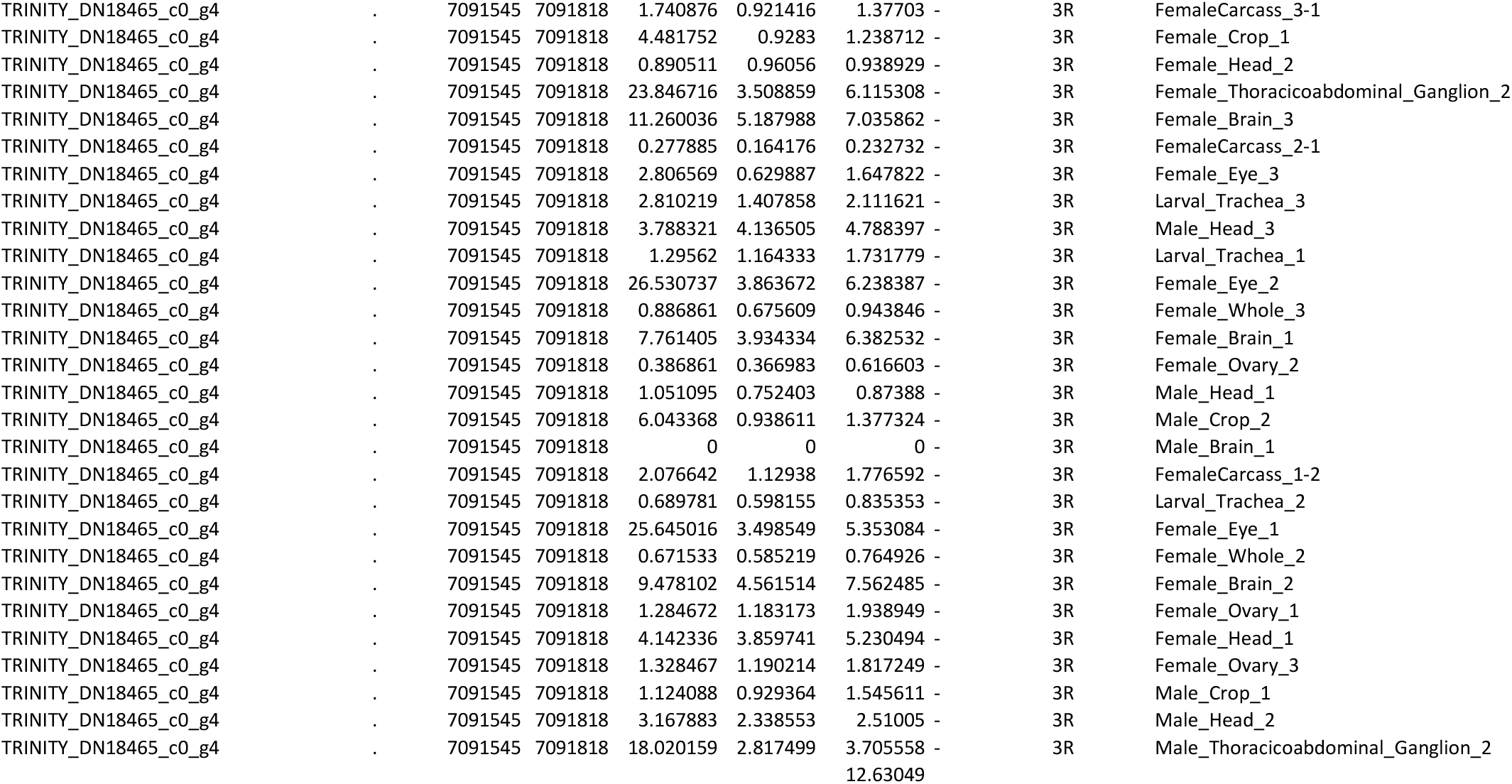
Expressed Candidate TPMs in FlyAtlas2 Libraries

**Supplemental Table 6.**
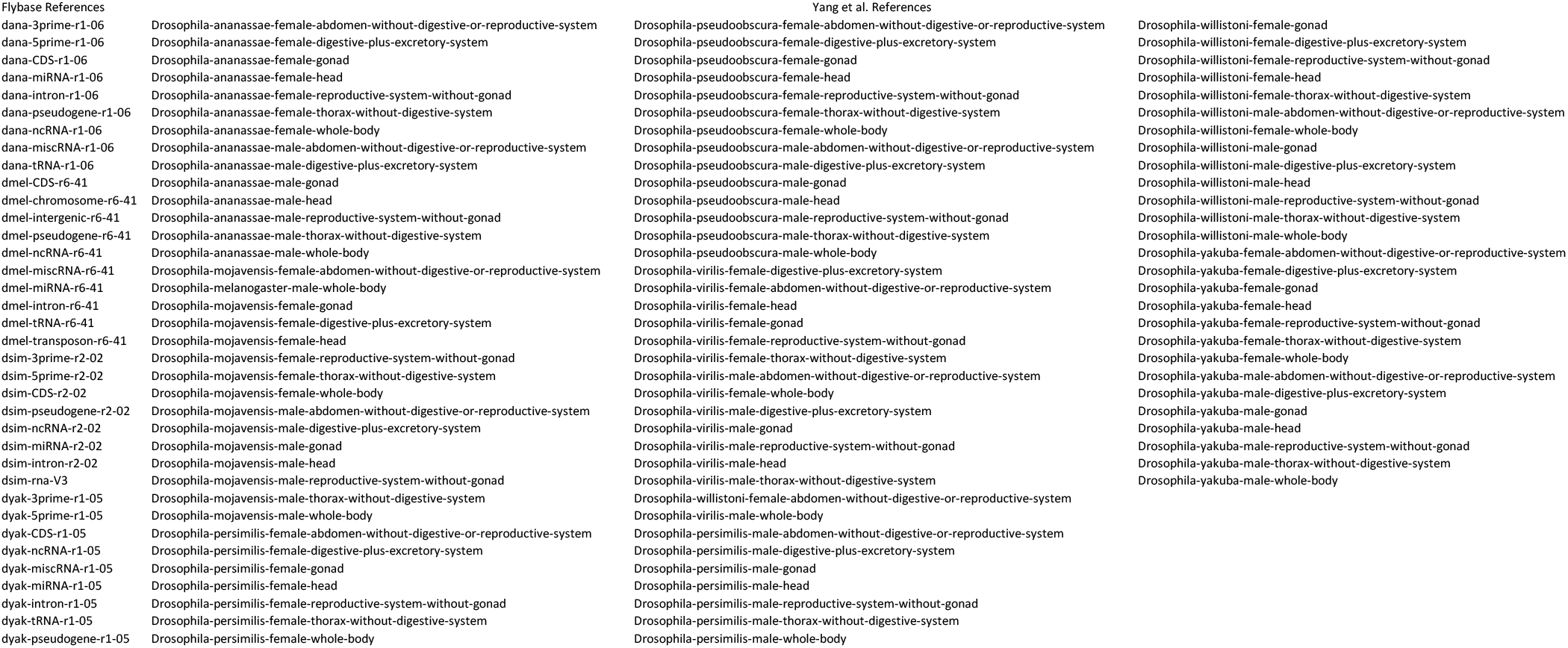
List of All BLAST Databases Utilized

## REFERENCES

Allen, A. K., & Spradling, A. C. (2008). The Sf1-related nuclear hormone receptor Hr39 regulates Drosophila female reproductive tract development and function. Development, 135(2), 311–321.

Baalsrud, H. T., Tørresen, O. K., Solbakken, M. H., Salzburger, W., Hanel, R., Jakobsen, K. S., & Jentoft, S. (2018). De novo Gene Evolution of Antifreeze Glycoproteins in Codfishes Revealed by Whole Genome Sequence Data. Molecular Biology and Evolution, 35(3), 593–606.

Begun, D. J., Lindfors, H. A., Kern, A. D., & Jones, C. D. (2007). Evidence for de novo evolution of testis-expressed genes in the Drosophila yakuba/Drosophila erecta clade. Genetics, 176(2), 1131–1137.

Begun, D. J., Lindfors, H. A., Thompson, M. E., & Holloway, A. K. (2006). Recently evolved genes identified from Drosophila yakuba and D. erecta accessory gland expressed sequence tags. Genetics, 172(3), 1675–1681.

Cai, J., Zhao, R., Jiang, H., & Wang, W. (2008). De novo origination of a new protein-coding gene in Saccharomyces cerevisiae. Genetics, 179(1), 487–496.

Carvunis, A.-R., Rolland, T., Wapinski, I., Calderwood, M. A., Yildirim, M. A., Simonis, N., Charloteaux, B., Hidalgo, C. A., Barbette, J., Santhanam, B., Brar, G. A., Weissman, J. S., Regev, A., Thierry-Mieg, N., Cusick, M. E., & Vidal, M. (2012). Proto-genes and de novo gene birth. Nature, 487(7407), 370–374.

Casola, C. (2018). From de novo to “de nono”: The majority of novel protein coding genes identified with phylostratigraphy are old genes or recent duplicates. In Genome Biology and Evolution. https://doi.org/10.1093/gbe/evy231

Cridland, J. M., Majane, A. C., Zhao, L., & Begun, D. J. (2022). Population biology of accessory gland-expressed de novo genes in Drosophila melanogaster. Genetics, 220(1). https://doi.org/10.1093/genetics/iyab207

Edgar, R., Domrachev, M., & Lash, A. E. (2002). Gene Expression Omnibus: NCBI gene expression and hybridization array data repository. Nucleic Acids Research, 30(1), 207–210.

Fowler, G. L. (1973). Some Aspects of the Reproductive Biology of Drosophila: Sperm Transfer, Sperm Storage, and Sperm Utilization. In E. W. Caspari (Ed.), Advances in Genetics (Vol. 17, pp. 293–360). Academic Press.

Grabherr, M. G., Haas, B. J., Yassour, M., Levin, J. Z., Thompson, D. A., Amit, I., Adiconis, X., Fan, L., Raychowdhury, R., Zeng, Q., Chen, Z., Mauceli, E., Hacohen, N., Gnirke, A., Rhind, N., di Palma, F., Birren, B. W., Nusbaum, C., Lindblad-Toh, K., … Regev, A. (2011). Full-length transcriptome assembly from RNA-Seq data without a reference genome. Nature Biotechnology, 29(7), 644–652.

Gramates, L. S., Agapite, J., Attrill, H., Calvi, B. R., Crosby, M. A., Dos Santos, G., Goodman, J. L., Goutte-Gattat, D., Jenkins, V. K., Kaufman, T., Larkin, A., Matthews, B. B., Millburn, G., Strelets, V. B., & the FlyBase Consortium. (2022). Fly Base: a guided tour of highlighted features. Genetics, 220(4). https://doi.org/10.1093/genetics/iyac035

Heames, B., Schmitz, J., & Bornberg-Bauer, E. (2020). A Continuum of Evolving De novo Genes Drives Protein-Coding Novelty in Drosophila. In Journal of Molecular Evolution (Vol. 88, Issue 4, pp. 382–398). https://doi.org/10.1007/s00239-020-09939-z

Heinen, T. J. A. J., Staubach, F., Häming, D., & Tautz, D. (2009). Emergence of a new gene from an intergenic region. Current Biology: CB, 19(18), 1527–1531.

Jin, G., Ma, P.-F., Wu, X., Gu, L., Long, M., Zhang, C., & Li, D.-Z. (2021). New Genes Interacted With Recent Whole-Genome Duplicates in the Fast Stem Growth of Bamboos. Molecular Biology and Evolution, 38(12), 5752–5768.

Kang, Y.-J., Yang, D.-C., Kong, L., Hou, M., Meng, Y.-Q., Wei, L., & Gao, G. (2017). CPC2: a fast and accurate coding potential calculator based on sequence intrinsic features. Nucleic Acids Research, 45(W1), W12–W16.

Karolchik, D., Baertsch, R., Diekhans, M., Furey, T. S., Hinrichs, A., Lu, Y. T., Roskin, K. M., Schwartz, M., Sugnet, C. W., Thomas, D. J., Weber, R. J., Haussler, D., Kent, W. J., & University of California Santa Cruz. (2003). The UCSC Genome Browser Database. Nucleic Acids Research, 31(1), 51–54.

Kim, D., Paggi, J. M., Park, C., Bennett, C., & Salzberg, S. L. (2019). Graph-based genome alignment and genotyping with HISAT2 and HISAT-genotype. Nature Biotechnology, 37(8), 907–915.

Krause, S. A., Overend, G., Dow, J. A. T., & Leader, D. P. (2022). FlyAtlas 2 in 2022: enhancements to the Drosophila melanogaster expression atlas. Nucleic Acids Research, 50(D1), D1010–D1015.

Leader, D. P., Krause, S. A., Pandit, A., Davies, S. A., & Dow, J. A. T. (2018). FlyAtlas 2: a new version of the Drosophila melanogaster expression atlas with RNA-Seq, miRNA-Seq and sex-specific data. Nucleic Acids Research, 46(D1), D809–D815.

Levine, M. T., Jones, C. D., Kern, A. D., Lindfors, H. A., & Begun, D. J. (2006). Novel genes derived from noncoding DNA in Drosophila melanogaster are frequently X-linked and exhibit testis-biased expression. Proceedings of the National Academy of Sciences of the United States of America, 103(26), 9935–9939.

Li, D., Dong, Y., Jiang, Y., Jiang, H., Cai, J., & Wang, W. (2010). A de novo originated gene depresses budding yeast mating pathway and is repressed by the protein encoded by its antisense strand. In Cell Research (Vol. 20, Issue 4, pp. 408–420). https://doi.org/10.1038/cr.2010.31

Li, H., Handsaker, B., Wysoker, A., Fennell, T., Ruan, J., Homer, N., Marth, G., Abecasis, G., Durbin, R., & 1000 Genome Project Data Processing Subgroup. (2009). The Sequence Alignment/Map format and SAMtools. Bioinformatics, 25(16), 2078–2079.

Long, M., Betrán, E., Thornton, K., & Wang, W. (2003). The origin of new genes: glimpses from the young and old. Nature Reviews. Genetics, 4(11), 865–875.

Luthra, I., Chen, X. E., Jensen, C., Rafi, A. M., Salaudeen, A. L., & de Boer, C. G. (2022). Biochemical activity is the default DNA state in eukaryotes. https://doi.org/10.1101/2022.12.16.520785

Mackay, T. F. C., Richards, S., Stone, E. A., Barbadilla, A., Ayroles, J. F., Zhu, D., Casillas, S., Han, Y., Magwire, M. M., Cridland, J. M., Richardson, M. F., Anholt, R. R. H., Barrón, M., Bess, C., Blankenburg, K. P., Carbone, M. A., Castellano, D., Chaboub, L., Duncan, L., … Gibbs, R. A. (2012). The Drosophila melanogaster Genetic Reference Panel. Nature, 482(7384), 173–178.

Manier, M. K., Belote, J. M., Berben, K. S., Novikov, D., Stuart, W. T., & Pitnick, S. (2010). Resolving mechanisms of competitive fertilization success in Drosophila melanogaster. Science, 328(5976), 354–357.

Mayhew, M. L., & Merritt, D. J. (2013). The morphogenesis of spermathecae and spermathecal glands in Drosophila melanogaster. Arthropod Structure & Development, 42(5), 385–393.

McDonough-Goldstein, C. E., Borziak, K., Pitnick, S., & Dorus, S. (2021). Drosophila female reproductive tract gene expression reveals coordinated mating responses and rapidly evolving tissue-specific genes. G3, 11(3). https://doi.org/10.1093/g3journal/jkab020

Murphy, D. N., & McLysaght, A. (2012). De novo origin of protein-coding genes in murine rodents. PloS One, 7(11), e48650.

Neme, R., & Tautz, D. (2013). Phylogenetic patterns of emergence of new genes support a model of frequent de novo evolution. BMC Genomics, 14, 117.

Palmieri, N., Kosiol, C., & Schlötterer, C. (2014). The life cycle of Drosophila orphan genes. eLife, 3. https://doi.org/10.7554/elife.01311

Pertea, M., Pertea, G. M., Antonescu, C. M., Chang, T.-C., Mendell, J. T., & Salzberg, S. L. (2015). StringTie enables improved reconstruction of a transcriptome from RNA-seq reads. Nature Biotechnology, 33(3), 290–295.

Pitnick, S., Marrow, T., & Spicer, G. S. (1999). EVOLUTION OF MULTIPLE KINDS OF FEMALE SPERM-STORAGE ORGANS IN DROSOPHILA. Evolution; International Journal of Organic Evolution, 53(6), 1804–1822.

Schnakenberg, S. L., Matias, W. R., & Siegal, M. L. (2011). Sperm-storage defects and live birth in Drosophila females lacking spermathecal secretory cells. PLoS Biology, 9(11), e1001192.

Sedghifar, A., Saelao, P., & Begun, D. J. (2016). Genomic Patterns of Geographic Differentiation in Drosophila simulans. Genetics, 202(3), 1229–1240.

Sun, J., & Spradling, A. C. (2012). Female Reproductive Glands Play Essential Roles in Reproduction That May Have Been Conserved During Evolution. Biology of Reproduction, 87(Suppl_1), 347–347.

Sun, J., & Spradling, A. C. (2013). Ovulation in Drosophila is controlled by secretory cells of the female reproductive tract. eLife, 2, e00415.

Teufel, F., Armenteros, J. J. A., Johansen, A. R., Gíslason, M. H., Pihl, S. I., Tsirigos, K. D., Winther, O., Brunak, S., von Heijne, G., & Nielsen, H. (2022). SignalP 6.0 predicts all five types of signal peptides using protein language models. In Nature Biotechnology (Vol. 40, Issue 7, pp. 1023–1025). https://doi.org/10.1038/s41587-021-01156-3

Thurmond, J., Goodman, J. L., Strelets, V. B., Attrill, H., Sian Gramates, L., Marygold, S. J., Matthews, B. B., Millburn, G., Antonazzo, G., Trovisco, V., Kaufman, T. C., Calvi, B. R., Perrimon, N., Gelbart, S. R., Agapite, J., Broll, K., Crosby, L., dos Santos, G., Emmert, D., … the FlyBase Consortium. (2019). FlyBase 2.0: the next generation. In Nucleic Acids Research (Vol. 47, Issue D1, pp. D759–D765). https://doi.org/10.1093/nar/gky1003

Vakirlis, N., Hebert, A. S., Opulente, D. A., Achaz, G., Hittinger, C. T., Fischer, G., Coon, J. J., & Lafontaine, I. (2018). A Molecular Portrait of De novo Genes in Yeasts. Molecular Biology and Evolution, 35(3), 631–645.

Van Oss SB, Carvunis A-R (2019) De novo gene birth. PLoS Genet 15(5): e1008160. https://doi.org/10.1371/journal.pgen.1008160

Wang, L., Park, H. J., Dasari, S., Wang, S., Kocher, J.-P., & Li, W. (2013). CPAT: Coding-Potential Assessment Tool using an alignment-free logistic regression model. Nucleic Acids Research, 41(6), e74.

Wilson, C., Leiblich, A., Goberdhan, D. C. I., & Hamdy, F. (2017). The Drosophila Accessory Gland as a Model for Prostate Cancer and Other Pathologies. In Current Topics in Developmental Biology (pp. 339–375). https://doi.org/10.1016/bs.ctdb.2016.06.001

Yang, H., Jaime, M., Polihronakis, M., Kanegawa, K., Markow, T., Kaneshiro, K., & Oliver, B. (2018). Re-annotation of eight genomes. Life Science Alliance, 1(6), e201800156.

Zhang, L., Ren, Y., Yang, T., Li, G., Chen, J., Gschwend, A. R., Yu, Y., Hou, G., Zi, J., Zhou, R., Wen, B., Zhang, J., Chougule, K., Wang, M., Copetti, D., Peng, Z., Zhang, C., Zhang, Y., Ouyang, Y., … Long, M. (2019). Rapid evolution of protein diversity by de novo origination in Oryza. Nature Ecology & Evolution, 3(4), 679–690.

Zhao, L., Saelao, P., Jones, C. D., & Begun, D. J. (2014). Origin and spread of de novo genes in Drosophila melanogaster populations. Science (New York, N.Y.), 343(6172), 769–772.

Zheng, E. B., & Zhao, L. (2022). Protein evidence of unannotated ORFs in reveals diversity in the evolution and properties of young proteins. eLife, 11. https://doi.org/10.7554/eLife.78772

Zhou Q., Meng Y., Su L., Zhao S.-M., Shi H.-P., & Huang S.-Z. (2008). [A Chinese girl with fibrodysplasia ossificans progressiva caused by a de novo mutation R206H in ACVR1 gene]. Zhonghua er ke za zhi. Chinese journal of pediatrics, 46(3), 215–219.

Zhou, Q., Zhang, G., Zhang, Y., Xu, S., Zhao, R., Zhan, Z., Li, X., Ding, Y., Yang, S., & Wang, W. (2008). On the origin of new genes in Drosophila. Genome Research, 18(9), 1446–1455.

Zhuang, X., & Cheng, C.-H. C. (2021). Propagation of a De novo Gene under Natural Selection: Antifreeze Glycoprotein Genes and Their Evolutionary History in Codfishes. Genes, 12(11). https://doi.org/10.3390/genes12111777

